# chromoMap: An R package for Interactive Visualization and Annotation of Chromosomes

**DOI:** 10.1101/605600

**Authors:** Lakshay Anand, Carlos M. Rodriguez Lopez

**Affiliations:** Environmental Epigenetics and Genetics Group, Department of Horticulture, College of Agriculture, Food and Environment, University of Kentucky, Lexington, Kentucky, USA

## Abstract

**Summary:** chromoMap is an R package for constructing interactive visualizations of chromosomes/chromosomal regions, and mapping of chromosomal elements (like genes) onto them, of any living organism. The package takes separate tab-delimited files (BED like) to specify the genomic co-ordinates of the chromosomes and the elements to annotate. Each rendered chromosome is composed of continuous loci of specific ranges where each locus, on hover, displays detailed information about the elements annotated within that locus range. By just tweaking parameters of a single function, users can generate a variety of plots that can either be saved as static image or shared as HTML documents. Users can utilize the various prominent features of chromoMap including, but not limited to, visualizing polyploidy, creating chromosome heatmaps, mapping groups of elements, adding hyperlinks to elements, multi-species chromosome visualization.

**Availability and implementation:** The R package *chromoMap* is available under the GPL-3 Open Source license. It is included with a vignette for comprehensive understanding of its various features, and is freely available from: https://CRAN.R-project.org/package=chromoMap.

**Supplementary information:** Supplementary data are available online.

## Introduction

The chromosomes store the information required to produce an entire living organism in its gigantic structural component, called the DNA, constituting sequence of four nucleotide bases. The dramatic decrease in the cost of genome sequencing, in recent years, has rapidly increased the amount of genome being sequenced and deposited to the public repositories. After sequencing, researchers are primarily interested to annotate chromosomal elements (like genes) to construct a physical map of the genome elucidating the structural boundaries of its annotated elements. Beside defining the structure, annotations can provide meaningful insights into understanding the genomic variations within and between species through genome comparisons.

Currently, genomes are usually viewed in interactive web-based genome browser applications like JBrowse[1]. There is, however, a paucity of non-web based independent tools capable of generating publication-ready interactive visualization and annotation of genomes. *ChromoMap*, developed as an R[2] package, is capable of generating interactive graphics visualization of entire chromosomes or chromosomal regions of any living organism facilitating annotation of thousands of chromosomal elements on a single plot. The humongous genome size of certain species, including our own, poses practical graphical challenges of displaying whole chromosomes within the available canvas space suitable for publications. The chromoMap, in order to surmount this restriction, render chromosomes as a continuous composition of loci. Each locus, consisting of a specific genomic range determined algorithmically based on chromosome length, displays the detailed information about the annotation in that region as a tooltip, thereby, imparting interactivity to the plot.

The chromosomes, and the annotations, are constructed by passing their genomic co-ordinates, and some optional secondary data, separately as tab-delimited text files similar to the commonly used bioinformatics file format, BED (Browser Extensible Data). A single R function, namesake of the package, is used for constructing a variety of different plots (discussed in the next section) by changing its various arguments values. The simplest annotation plot, however, can be constructed by using the following command:

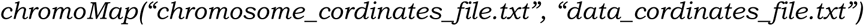

The first and second arguments, respectively, taking the co-ordinates of the chromosomes and the annotations. The generated plot, as viewed in RStudio’s viewer pane, offers interactivity to the users to view deeply into a locus allowing them to explore their annotations of interest. Users can export the plot either as a static image or as a stand-alone HTML file encompassing the interactive plot capable of including as supplementary data in publications. In addition, the plots can be included in Shiny applications, or embedded in RMarkdown documents. The rest of the paper illuminates the user about some of the most prominent features offered by the package illustrating, with examples. its application in research field.

## Salient features and applications

### Polyploidy

Inferable from its name, the polyploidy feature allows to construct sets of chromosomes simultaneously on the same plot. This is achieved by simply passing co-ordinates files separately for each set of chromosomes and setting the argument *ploidy* equivalent to the number of sets being passed. Each set of chromosomes are rendered independent of each other and, hence, can take sets of chromosomes that differ in size and numbers. An interesting plausible application of this feature is to visualize cross-species chromosome sets that can be advantageous for their comparative studies. Figure 1 describes the use of this feature to annotate orthologous genes in rice and maize conspicuously depicting the notable difference in their genome sizes and chromosome numbers. Polyploidy visualizations can be achieved by using the following command:

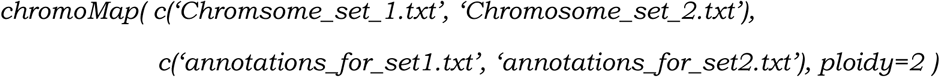

**Figure 1.**
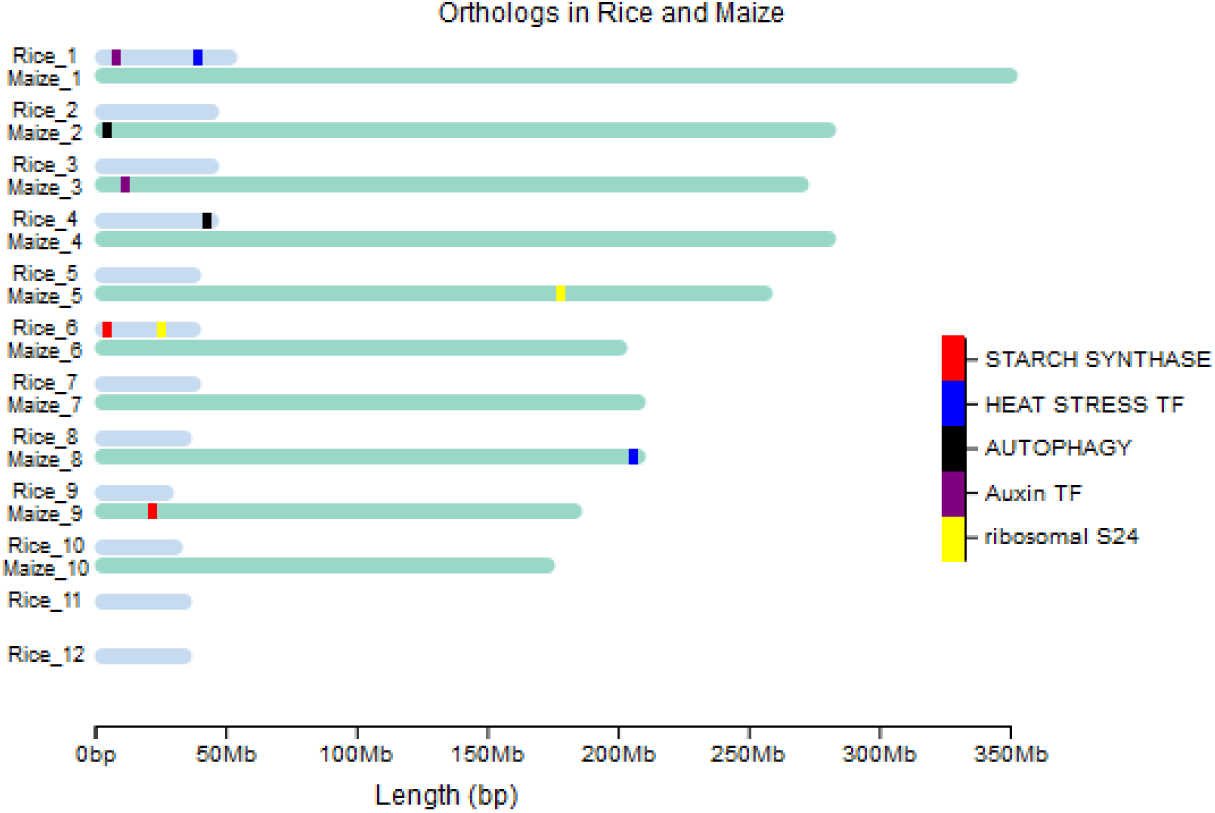
Visualizing orthologs in rice and maize using *chromoMap*. Five randomly selected sets of orthologous genes, shown as distinct colors, were fetched from Ensembl Plants database[3]. The interactive version of the plot (supplied in supplementary files) shows each gene identifier name on hover. Each locus, in this case, shows a single gene since it’s the only gene annotated in that locus range.

### Group annotations

Many instances of visualizing annotations, including the one shown in figure 1, require depicting annotations of groups of elements as opposed to individual ones. To visually differentiate between groups, chromoMap allows users to assign groups, for each element being passed, as an additional data column in input file. Figure 2 depicts another example of polyploidy and group annotation where the paralogous and homeologous genes in allopolyploid wheat genome are visualized.

**Figure 2.**
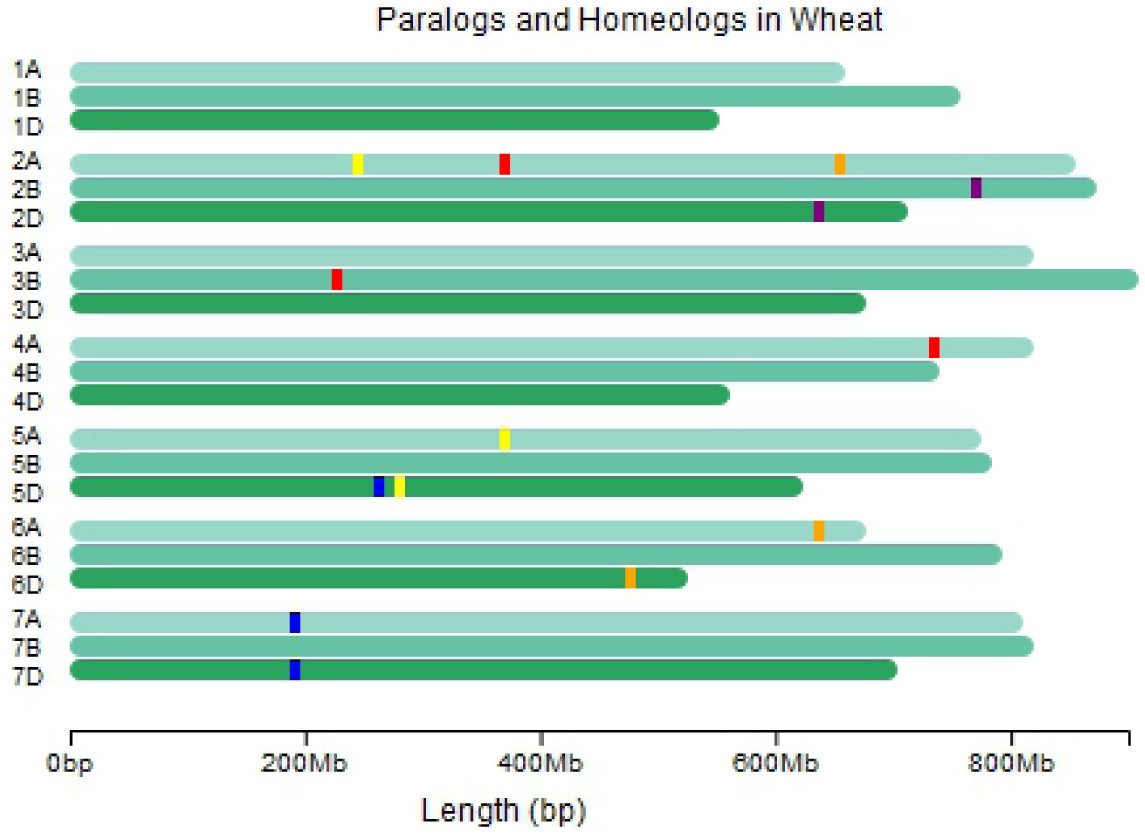
Visualization of allopolyploid wheat genome delineating the paralogous and homeologous genes created using *chromoMap*. Five gene sets are shown in distinct colors.

### Point and Segment-annotations

For annotating elements on the chromosomes, ChromoMap provides the choice of two annotation algorithms, viz. point-annotation and segment-annotation, differing in how annotations are visualized on the plot. The point annotation algorithm ignores the size of the element, and, annotate the element on a single locus. The segment annotation algorithm, on the other hand, consider the size of the element visualizing the annotation as a segment. This can be advantageous when visualizing chromosomal regions, like gene, annotating its structural elements (figure 3).

**Figure 3.**
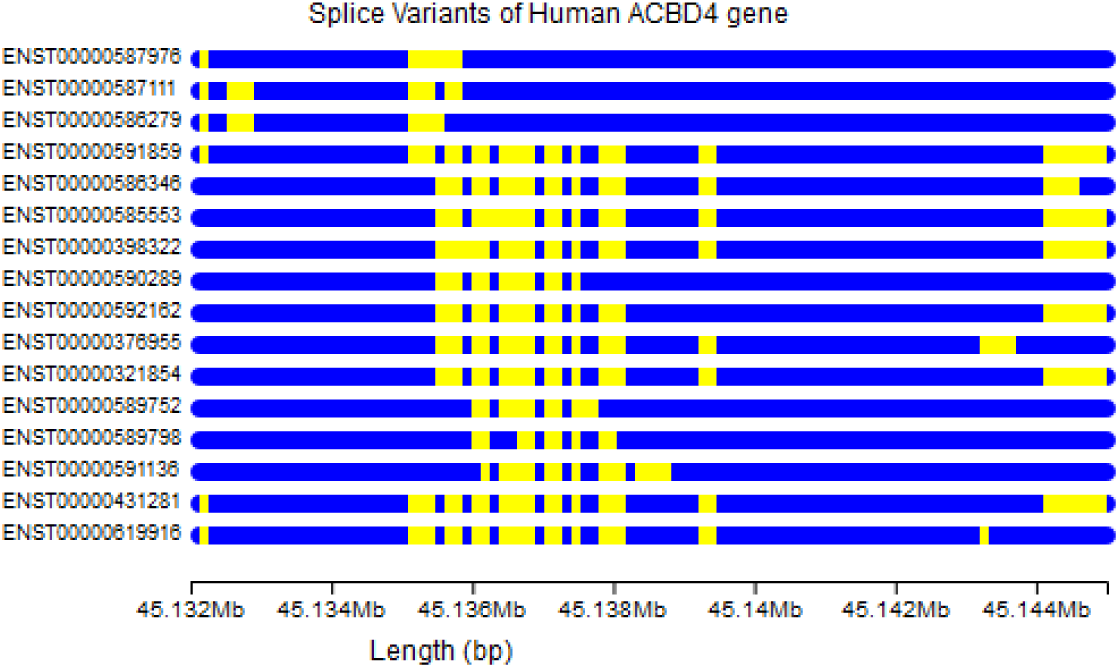
Visualizing splice variants in human gene ACBD4. Genomic co-ordinates were fetched from Ensembl.

### Chromosome heatmaps

The element-associated numeric data, such as gene expression values, can be visualized effectively as heatmaps. As each locus represents a specific range of base pairs, multiple elements can possibly be annotated in its range. chromoMap uses a data aggregation algorithm, provided as average or sum functions, to assign data values to each locus. Hence, for the locus encompassing more than one elements, aggregate data value will be assigned to it. The actual data values, however, can still be viewed in tooltip for each of the element. The differential expression analysis results from a breast cancer study from GEO(Gene Expression Omnibus)[4], shown in figure 4, utilizes this feature in combination of other features of chromoMap. The chromosome strands show the average expression change(logFC) per loci, while the mean expression (normalized) values from the two groups (control and case) has been shown in adjacent.

**Figure 4.**
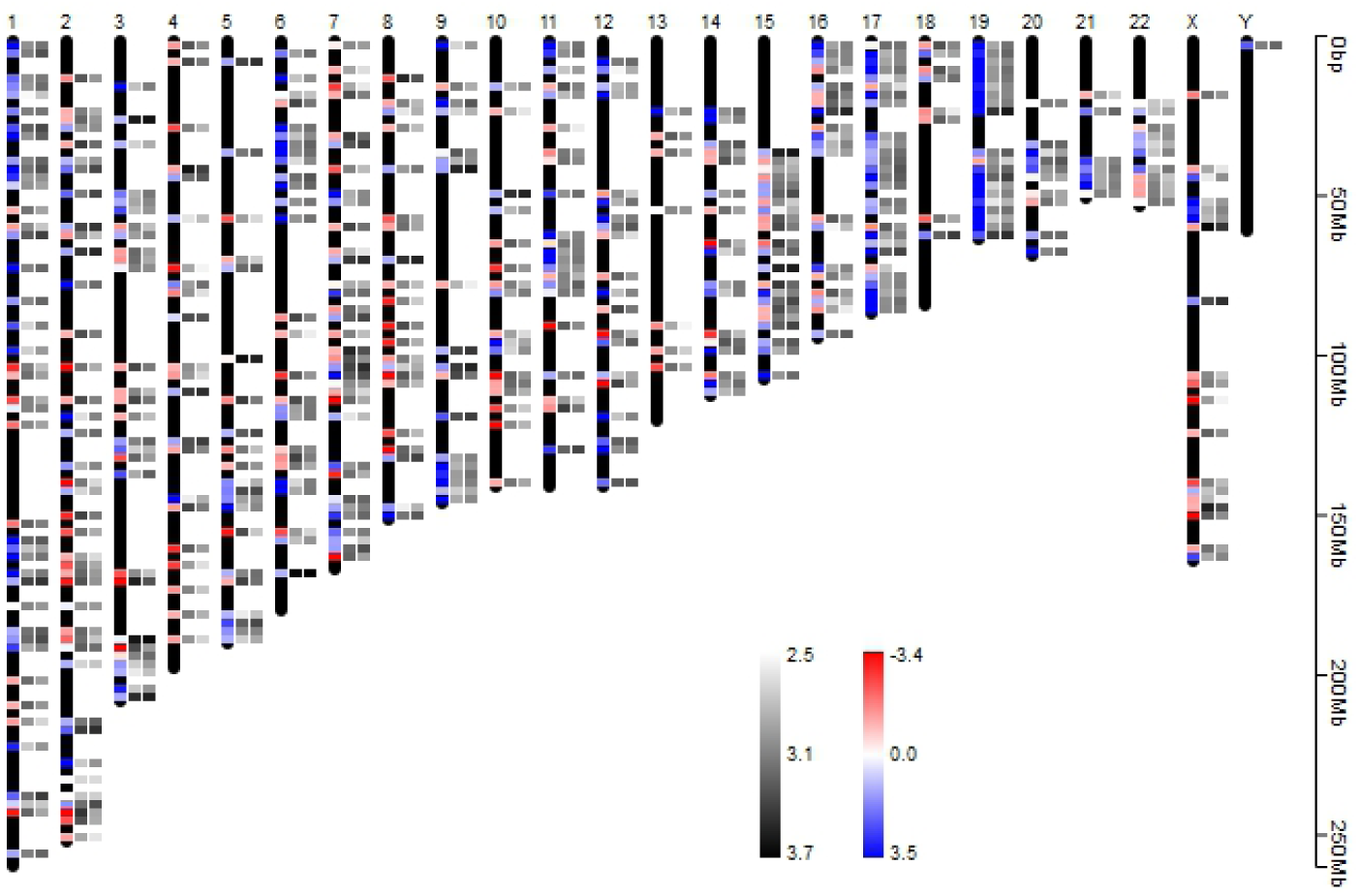
Visualizing differential expression analysis results from a breast cancer study data[5]. Chromosomes in black strand depict average expression change(logFC) Per loci where red and blue regions signify down-regulation and up-regulation respectively. The mean of normalized expression of the two groups, for the same genes, is shown in adjacent (control and case).

### Hyperlinks and labelling

While interactivity allow the users to lookup in detail about the annotations, the chromoMap also provide the feature of associating hyperlinks for each annotation on the plot. Hence, when the genes are viewed in tooltips, users can directly click on them and land to the web page describing its details. Moreover, the users can use the labelling option to display labels on the plot generating a static visualization of the plot.

## Supporting information

Interactive Figures

## Acknowledgement

All the data required for constructing plots and annotations were fetched from public repositories like UCSC, Ensembl, and NCBI.

